# NCLX prevents cell death during adrenergic activation of the brown adipose tissue

**DOI:** 10.1101/464339

**Authors:** Essam A. Assali, Anthony E. Jones, Michaela Veliova, Mahmoud Taha, Nathanael Miller, Michaël Shum, Marcus F. Oliveira, Guy Las, Marc Liesa, Israel Sekler, Orian S. Shirihai

## Abstract

A sharp increase in mitochondrial Ca^2+^ marks the activation of the brown adipose tissue (BAT) thermogenesis, yet the mechanisms preventing Ca^2+^ deleterious effects are poorly understood. Here, we show that adrenergic stimulation of BAT activates a PKA-dependent mitochondrial Ca^2+^ extrusion via the mitochondrial Na^+^/Ca^2+^ exchanger, NCLX. Adrenergic stimulation of NCLX-ablated brown adipocytes (BA) induces a profound mitochondrial Ca^2+^ overload and impaired uncoupled respiration. Core body temperature, PET imaging and VO2 measurements confirm a BAT specific thermogenic defect in NCLX-null mice.

We show that mitochondrial Ca^2+^ overload induced by adrenergic stimulation of NCLX-null BAT, triggers the opening of the mitochondrial permeability transition pore (mPTP), leading to remarkable mitochondrial swelling, Cytochrome *c* release and cell death in BAT. However, treatment with mPTP inhibitors rescue mitochondrial respiratory function and thermogenesis in NCLX-null BA, *in vitro* and *in vivo*.

Our findings identify a novel pathway enabling non-lethal mitochondrial Ca^2+^ elevation during adrenergic stimulation of uncoupled respiration. Deletion of NCLX transforms the adrenergic pathway responsible for the stimulation of thermogenesis into a death pathway.

## Introduction

The brown adipose tissue (BAT) dissipates energy in the form of heat after hormonal or adrenergic stimuli triggered by cold-exposure^1,2^. This pathway is initiated by the sympathetic neurotransmitter Norepinephrine (NE), which induces a PKA-dependent lipid mobilization and oxidization. Resultant free-fatty acids (FFAs) activate BAT signature protein, Uncoupler Protein 1 (UCP1) leading to proton leak and uncoupled respiration ^1,3,4^. In addition to their role in inducing UCP-1 function, FFAs undergo β-oxidation and fuel the TCA cycle for production of reducing equivalents, to supply the electron transport chain (ETC), essentially utilized to drive uncoupled energy expenditure^2^. However, for TCA cycle to meet up with the energy demand of thermogenesis, mitochondrial Ca^2+^ is required for upregulating the activity of the dehydrogenases and ETC components^5–7^. Yet, uncontrolled or prolonged mitochondrial Ca^2+^ rise, often encountered during ischemic^8^ or neurodegenerative diseases^9^, can lead to opening of the mitochondrial permeability transition pore (mPTP)^10,11^.

The opening of mPTP has been associated with two hallmarks, the first is depolarization and a collapse in membrane potential, which is accompanied by excess of mitochondrial Ca^2+^ accumulation^8,12^. The association between mitochondrial Ca^2+^ overload and depolarization is not well-understood, yet both stimuli were shown to be essential and sufficient for the pore opening, leading to mitochondrial swelling and triggering Cytochrome *c* release; thus impairing respiration and culminating in cellular death^12–14^. Thereby, the fine-tuning of mitochondrial Ca^2+^ homeostasis in conjugation with thermogenesis, has to be tight and well-regulated to avoid detrimental effects on cell physiology.

Mitochondrial Ca^2+^ uptake is mediated through a highly selective Ca^2+^ channel, recently linked to the mitochondrial calcium uniporter (MCU) gene ^15,16^, Ca^2+^ is subsequently transported out primarily by a mitochondrial Na^+^/Ca^2+^ exchanger termed NCLX^17^. These components were known for decades but their molecular identities have been discovered only recently^18^.

The contribution of mitochondrial Ca^2+^ signaling to thermogenesis upon cold-stress is poorly understood and little is known about its role in BAT, despite its potential importance based on early studies, where it was shown that Na^+-^dependent mitochondrial Ca^2+^ extrusion is essential for brown adipocyte (BA) uncoupled-respiration^19^. However, the involvement of this process in BAT thermoregulation or its mode of regulation remained largely unknown.

A recent study showed that that PKA-mediated phosphorylation of NCLX, at a serine residue (Ser258) located on the regulatory site of the protein rescued mitochondrial Ca^2+^efflux in depolarized neurons lacking PINK1^20^. However, the physiological role of this PKA-dependent regulation is still unclear.

In this study, we show that adrenergic signaling activates a NCLX-mediated mitochondrial Ca^2+^ efflux required to establish both uncoupled energy expenditure and BAT cellular viability.

In the absence of adrenergic stimulation, deletion of the NCLX is inconsequential. While adrenergic stimulation of WT BA elicits thermogenic response characterized by a robust increase in mitochondrial respiration, adrenergic stimulation of BA lacking NCLX results in Ca^2+^ overload, mitochondrial swelling and Cytochrome *c* release leading to cell death in BAT. We found that mPTP inhibition fully restores the thermogenic capacity of NCLX-null BA both *in vitro* and *in vivo*. Overall, this study reveals a novel pathway through which mitochondrial Ca^2+^ efflux permits a robust activation of respiration in response to profound uncoupling, while preventing the activation of cell death pathways.

## Results

### Mitochondrial Ca^2+^extrusion is regulated by PKA activity in BAT

Cell stimuli mediated by different signaling molecules and hormones prompt an increase of cytosolic Ca^2+^ signals either from the influx of extracellular Ca^2+^ via the plasma membrane Ca^2+^ channels or the release of Ca^2+^ from internal stores, mostly via the 1,4,5-triphosphate receptor (IP3R)^21^. Increase of cytosolic Ca^2+^ leads to an increase in its uptake by the mitochondria and is succeeded by its extrusion from the organelle as demonstrated in Figure 1a upon NE stimulation. Noticeably, Ca^2+^ influx rate is faster than its efflux rate, indicating that the Ca^2+^ extrusion entity is the rate-limiting controller of mitochondrial Ca^2+^ transients due to its slower activity **(Fig. 1a)**.

**Figure 1.**
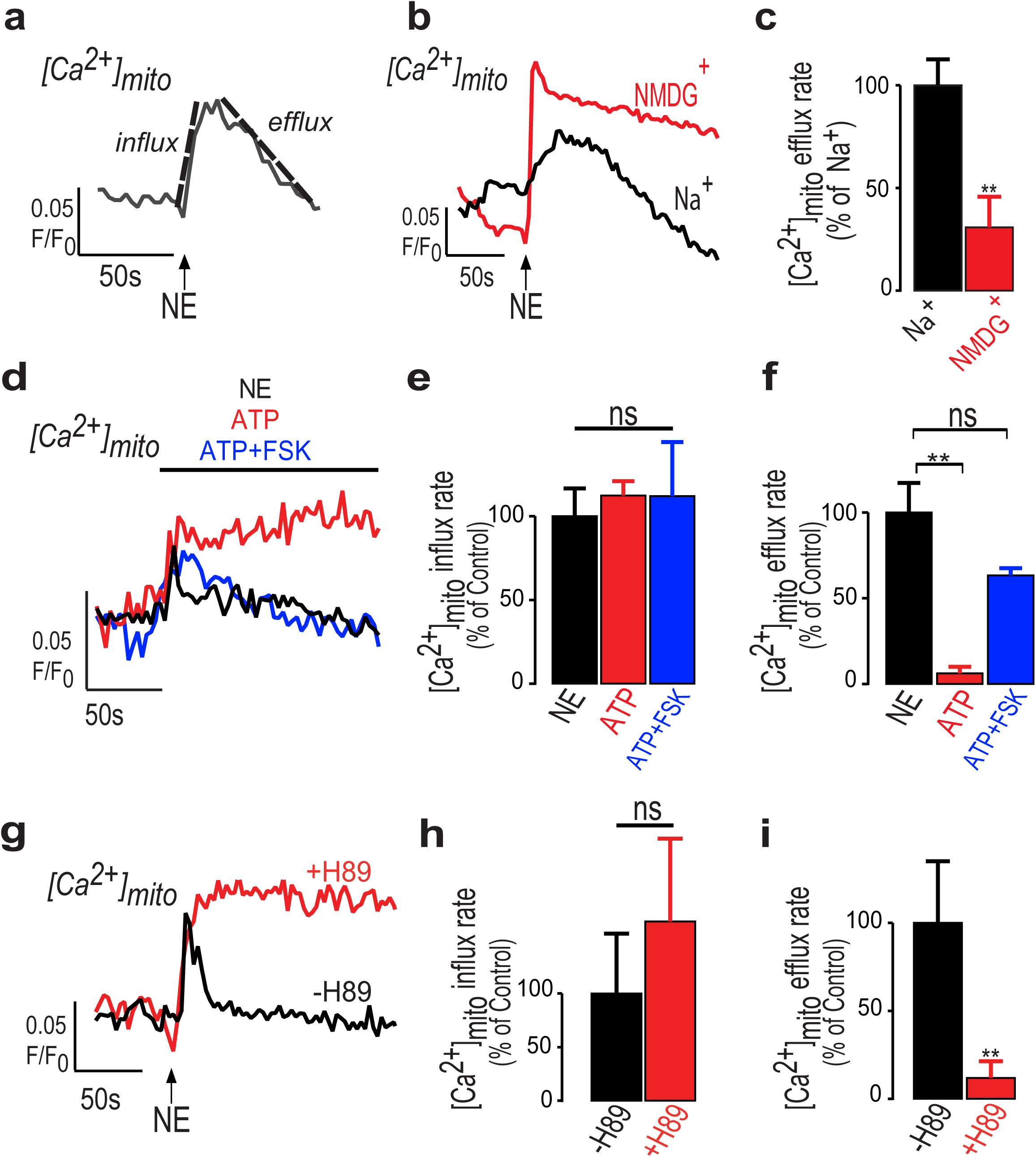
Mitochondrial Ca^2+^ extrusion in BA is mediated by Na^+^/Ca^2+^ exchange and requires PKA. **a:** Representative fluorescent traces of mitochondrial Ca^2+^ kinetics in BA after application of NE (1.5 μM). Mitochondrial Ca^2+^ is monitored using the mitochondrial Ca^2+^ dye, Rhode2-AM. The dashed lines represent the linear fit used to calculate the Ca^2+^ influx and efflux rates. **(b,c):** NE mediates a Na^+^-dependent mitochondrial Ca^2+^ efflux in primary BA. **b:** Representative fluorescent traces of mitochondrial Ca^2+^ transients upon application of NE to BA in the presence or absence of Na^+^ (NMDG^+^ iso-osmotically replacing Na^+^). Absence of Na+ strongly reduced mitochondrial Ca^2+^ efflux indicating that the mitochondrial Na^+^/Ca^2+^exchanger is the main mitochondrial Ca^2+^ exporter in BAT. **c:** Quantification of mitochondrial Ca^2+^ efflux rates in the presence and absence of Na^+^. NMDG^+^ is used as a cationic replacement of Na^+^ (n= 4-5 independent experiments per condition). **(d-f):** PKA activity is required for mitochondrial Ca^2+^ efflux in primary BA. **d:** Representative fluorescent traces of mitochondrial Ca^2+^ transients upon application of NE (1.5 μM), ATP (100μM) or ATP under pretreated cells with PKA activator, Forskolin (FSK (50 μM, 15 min)). Note that ATP induces a PKA–independent transient increase in mitochondrial Ca^2+^ that is not followed by an efflux. **e,f:** Quantification of mitochondrial Ca^2+^ fluxes in response to NE, ATP or ATP+FSK stimulation. Note that none of the Ca^2+^stimulants affected mitochondrial Ca^2+^ influx rate. However, Ca^2+^efflux was suppressed under ATP stimulation. FSK pretreatment of ATP stimulated cells completely restored Ca^2+^efflux. NE (n = 12), ATP (n = 4), ATP+FSK (n = 6). **(g-i):** PKA inhibition blunts Ca^2+^ efflux under NE treatment. **g:** Representative fluorescent traces of mitochondrial Ca^2+^ transients in primary BA upon stimulation by NE in the presence and absence of PKA inhibitor, H-89 (5μM, 1 h pre-incubation). **h,i:** Quantification of mitochondrial Ca^2+^ traces under treatment of H-89. Note that H-89 inhibits Ca^2+^ efflux while sparing Ca^2+^ influx, indicating that NCLX regulation by PKA is essential for NE-dependent mitochondrial Ca^2+^ efflux in BA. (n = 8 – 14 independent experiments per condition). Student’s t-test (**c,h,i**); One-way ANOVA with Tukey’s post-hoc test (**e,f**). Data are expressed as means ± SEM. ns p > 0.05, *p < 0.05, ** p < 0.001.

We initially determined whether Ca^2+^ efflux is Na^+^-dependent in intact BA upon physiological stimulation by superfusing primary BA cells with NE while monitoring mitochondrial Ca^2+^ fluxes using the mitochondrial Ca^2+^ dye, Rhod-2 AM. We found that mitochondrial Ca^2+^ release, but not its uptake, is dependent on Na^+^ in BA. Substituting Na^+^ iso-osmotically with NMDG^+^ resulted in a strong reduction in mitochondrial Ca^2+^ efflux. Therefore, we conclude that the mitochondrial Na^+^/Ca^2+^ exchanger is an essential mechanism for mitochondrial Ca^2+^extrusion in BAT **(Fig. 1b,c)**.

Although NE is the physiological stimulus of BAT, this agonist has a broad activity besides inducing intracellular Ca^2+^ rise. Therefore, we applied the purinergic agonist (ATP) to induce intracellular Ca^2+^ transients similar to the previous system. ATP, a broadly-studied purinergic agonist mobilizes Ca^2+^ to the cytosol and subsequently to the mitochondria by depleting the ER stores via IP3R^22^. Unlike NE, ATP activity does not involve stimulation of Gs alpha subunit and induction of PKA signaling^22–24^. Unexpectedly however, Ca^2+^ efflux, but not Ca^2+^ uptake, was almost totally inhibited when the cells were stimulated by ATP as compared to NE. Yet, this inhibition was reversed when the cells were pretreated with the PKA agonist, Forskolin (FSK), indicating that under physiological activity of BAT, NE signaling is amplifying a PKA-mediated pathway that is required to upregulate mitochondrial Ca^2+^ efflux **(Fig. 1d-f)**.

To further validate the role of PKA signaling in the activation of mitochondrial Ca^2+^ efflux, we stimulated BA with NE in the presence or absence of the PKA inhibitor, H-89 and mitochondrial Ca^2+^ was monitored. Consistent with the results presented in Figure 1d-f, mitochondrial Ca^2+^ efflux, but not the influx, was entirely blocked by H-89, further supporting that PKA signaling is essential for regulating mitochondrial Ca^2+^ efflux in BAT **(Fig. 1g-i)**.

Overall, these data show that NE regulates mitochondrial Ca^2+^efflux, but not influx, in a PKA-dependent manner.

### NCLX activity is essential for NE-induced mitochondrial Ca^2+^ efflux and uncoupled respiration in BA

Next, we asked whether NCLX is essential for mitochondrial Ca^2+^ efflux in activated BA and then tested its role in uncoupled respiration and energy-expenditure. First, mitochondrial Ca^2+^ signaling was monitored in primary BA transfected with small interfering RNA, siNCLX or siControl **(Fig. 2a-c)**; and in isolated BA from NCLX-Knock-Out (KO) and WT mice **(Fig. 2di)**. In both experimental paradigms, mitochondrial Ca^2+^ efflux was totally inhibited **(Fig. 2c,i)**, promoting a sustained mitochondrial Ca^2+^ overload after adrenergic stimulation. Our results further show that adrenergic stimulation did not affect mitochondrial Ca^2+^ uptake **(Fig. 2b,h)** or result in alterations of MCU expression levels tested in NCLX-ablated BA **(Fig. 2e,f)**. Collectively, these data indicate that NCLX is the primary Ca^2+^exporter in BAT mitochondria.

**Figure 2:**
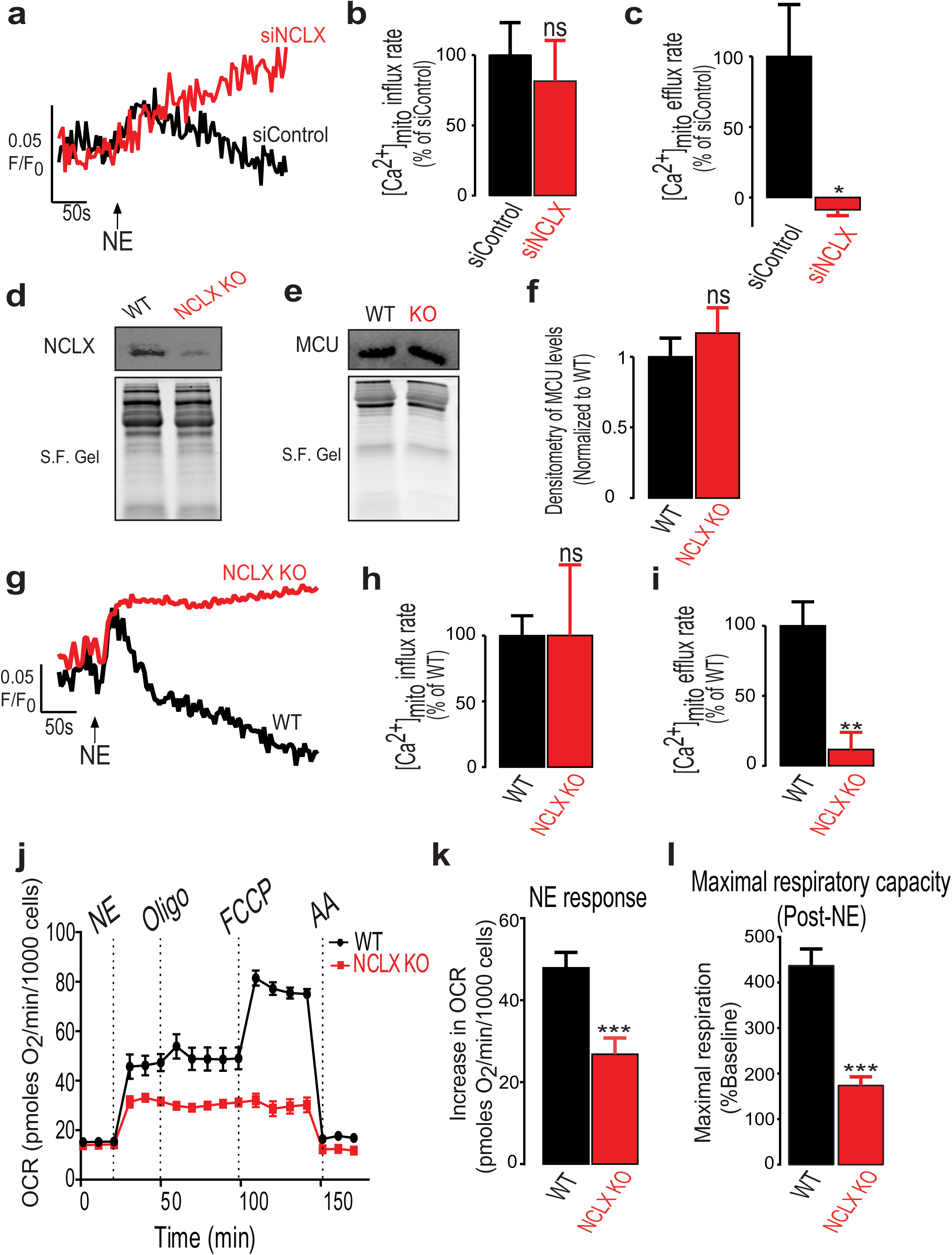
NCLX is required for mitochondrial uncoupled respiration in primary BA. **(a-c):** NCLX silencing promotes mitochondrial Ca^2+^ overload upon NE stimulation. **a:** Representative fluorescent traces of mitochondrial Ca^2+^ fluxes in BA transfected either with siNCLX or siControl and monitored for mitochondrial Ca^2+^ transients evoked by NE. **b,c:** Quantification of influx and efflux rates in primary BA transfected either with siNCLX or siControl. Downregulation of NCLX inhibited Ca^2+^ efflux while leaving Ca^2+^ influx unaffected (n = 4 - 5 independent experiments per condition). **d:** Western blot analysis of NCLX levels from NCLX KO and WT BA. Stain-Free Gels (S.F. Gels) are used as a normalization method for total loaded protein. **e,f:** Western blot analysis of MCU and quantification of its relative levels. MCU levels were not changed in BA of NCLX KO mice as compared to WT. (n=11 per group). **(g-i):** Adrenergic stimulation leads to mitochondrial Ca^2+^ overload in NCLX KO BA. **g:** Representative fluorescent traces of BA from NCLX KO and WT mice monitored for mitochondrial Ca^2+^ evoked by NE. **h,i:** Quantification of Ca^2+^ influx and efflux rates in NCLX KO and WT BA. mitochondrial Ca^2+^influx rate was not altered while Ca^2+^efflux was blocked. (n=4-12 experiments per condition). **(j-l):** Respiration measurements to address loss of NCLX effect on mitochondrial bioenergetics. **j:** Representative traces of oxygen consumption rates (OCR) of cultured BA from NCLX KO and WT mice. Stimulation by NE was the first injection. OLIGOmycin was used to assess mitochondrial uncoupling efficiency. FCCP was then injected to assess maximal respiration and non-mitochondrial OCR was evaluated by Antimycin A injection (AA). Note that NCLX KO BA have reduced adrenergic stimulated OCR and lower maximal respiration. **k,l:** Quantification of NE response and maximal respiration after NE stimulation. Note that NE-induced uncoupled respiration is suppressed in NCLX KO BA, as well as the loss of maximal respiration demonstrated by the inability of FCCP to further uncouple the mitochondria, indicating that NCLX loss impairs mitochondrial electron transport chain activity in stimulated BA. (N=3 independent experiments with n=18-22 wells per condition). Student’s t-test (**b,c,f,h,i,k,l**); Data are expressed as means ± SEM. ns p > 0.05, *p < 0.05, ** p < 0.001, *** p < 0.0001.

Subsequently, we investigated the bioenergetic consequences of NCLX deletion in primary BA. We hypothesized that NCLX activity will be required to maintain NE-stimulated mitochondrial uncoupling and energy expenditure in BA. To test this hypothesis, we compared NE-stimulated oxygen consumption rates of primary BA from NCLX KO mice and WT mice. As shown in Figure 2j, the increase in oxygen consumption rate (OCR) induced by NE is a measure of mitochondrial uncoupled respiration in BA, at physiological conditions. This is validated by the negligible effect of the ATP synthase inhibitor, oligomycin, on NE stimulated respiration (Oligomycin injection, Fig. 2j).

Our results show that that uncoupled respiration due to adrenergic stimulation by NE is remarkably suppressed in NCLX KO BA, losing close to half of the capacity as compared to WT group **(Fig. 2j,k)**. To determine if reduced energy expenditure was a result of reduced respiratory capacity or decreased activation of proton leak by UCP1 we employed an artificial uncoupler, FCCP. Maximal capacity for uncoupled respiration was tested in BA by injection of FCCP into the respirometer chamber after NE. Our results show that adrenergically stimulated NCLX-null BA have a strong decrease in their mitochondrial maximal respiration compared to stimulated WT, with a maximum of 437.4 ± 36.64% of basal respiration for WT compared to 173.8 ± 19.37% for NCLX-null BA **(Fig. 2j,l)**.

We further recapitulated this system in primary BA from WT mice, and subjected the cells to a Na^+^-containing medium or a Na^+^-depleted medium, by iso-osmotically substituting Na^+^ with NMDG^+^. Similar to the previous bioenergetics experiment, we observed that removal of Na^+^ caused a reduction in adrenergic respiratory response, followed by suppression of maximal respiration **(Supplementary Fig. 1a-d)**. Unlike the NCLX-null BA however, basal respiration was reduced in this experimental system, which can be attributed to off-target effect of lack of Na^+^ on metabolic functions **(Supplementary Fig. 1a)**.

In all, we show that the NCLX is a critical component for the regulation of BA respiratory rates and thermogenesis, induced by adrenergic stimulation, to maintain mitochondrial uncoupling and prevent Ca^2+^ overload.

### NCLX is essential for BAT thermogenic function in vivo

Based on the observed impairment in adrenergic-stimulated uncoupled respiration of NCLX-deficient BA *in vitro*, we assessed the role of NCLX in BAT thermogenesis by comparing in NCLX KO mice to their WT.

We used several complementary approaches to evaluate BAT energy expenditure and thermogenic capacity. First, we determined defense of body temperature after acute response to cold stress by continuously measuring mice core body temperature at ambient 4°C. Our results show a major impairment of body temperature defense in NCLX KO male mice after acute cold exposure for 6-8 h, as nearly 70% of the mice were pulled out due to their impaired thermogenic function. This is depicted in the survival curve where the term *survival* indicates the capacity to maintain body temperature above 28°C **(Fig. 3a,b)**. This finding was further confirmed in another cohort of NCLX KO and WT male littermates **(Supplementary Fig. 2a,b)**. We then monitored mice whole body oxygen consumption during cold stress at 4°C. No difference in the mice oxygen consumption was observed at time=0. However, at T=7 h whole body oxygen consumption was significantly reduced in NCLX KO compared to WT mice, with substantial loss of more than 40% of their O_2_ consumption **(Fig. 3c)**.

**Figure 3:**
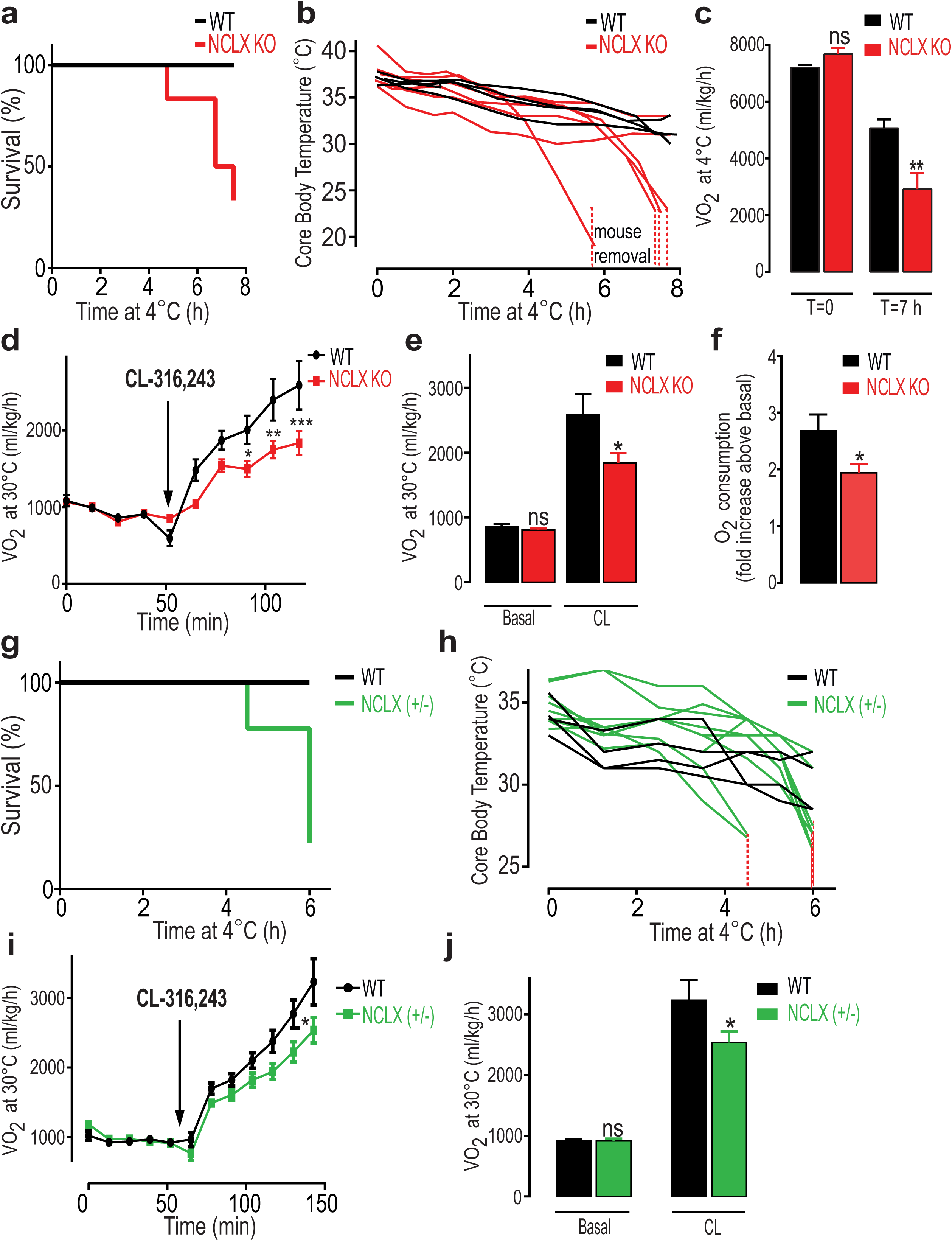
Thermogenesis is impaired in NCLX-null mice. **(a,b):** Cold tolerance tests of NCLX KO and WT male mice at 4°C with their raw core body temperature traces and survival curves (n =4-6 mice per group). **a:** Survival curves of 8-10 weeks old NCLX KO and WT male mice cold-stressed at 4°C. Animals reaching temperatures 28°C and below were returned to room temperature for recovery and counted as drop-outs. **b:** Core body temperature traces of the animals from experiment (a) during cold exposure (4°C). Note that the NCLX KO animals display impaired thermogenic capacity compared to their WT. Dashed red line indicates animal removal. **c:** VO2 of NCLX KO and WT mice subjected to cold (4°C) at t=0 and t=7h. Note that *O_2_* consumption was similar for both groups at t=0. However, NCLX KO mice lost their respiratory capacity at t=7 h compared to the WT (n=5 mice per group). **(d-f)**: Non-shivering thermogenesis of NCLX KO and WT mice was evaluated by measuring *O_2_* consumption at baseline and followed by β-3 adrenergic stimulation (CL-316,243 injection (1 mg/kg)) in anesthetized mice at 30°C (n= 5-6 mice per group). **d:** Traces of VO2 of NCLX KO and WT mice. **e:** VO2 at baseline and under CL-stimulation. Note that NCLX KO mice have reduced ability to increase their BAT stimulated *O_2_* consumption compared to WT. **f:** Fold increase of *O_2_* consumption after the CL-stimulation. (g-h) Survival curves of cold-stressed WT and NCLX (+/-) mice with their raw core body temperature (n= 4-9 mice per group). **g:** Survival curves of WT and NCLX (+/-) littermates cold-stressed at 4°C, Mice reaching 28°C or lower were returned to room temperature for recovery and counted as drop-outs. **h:** Core body temperature of the animals from the same experiment in (g). Note that NCLX (+/-) mice too, have an impaired thermogenic capacity as compared to WT. **(i,j):** VO2 of WT and NCLX (+/-) mice at basal and after CL316, 243-injection (1 mg/kg), under anesthesia at 30°C. (n = 4 mice per group). **i:** Traces of VO2 of NCLX (+/-) and WT mice. **j:** VO2 at baseline and under CL-stimulation. Note that NCLX (+/-) mice have reduced ability to increase their BAT stimulated *O_2_* consumption compared to WT. Student’s t-test (**c,e,f,j**); Two-way ANOVA (**d,i**). Data are expressed as means ± SEM. ns p > 0.05, *p < 0.05, ** p < 0.001, *** p < 0.0001.

To further dissect BAT non-shivering thermogenesis from shivering components controlled by the muscles, we measured oxygen consumption in response to a selective β3-adrenergic agonist, CL-326,243 in anesthetized mice. CL-326,243 was injected to NCLX KO and WT mice that were kept under thermoneutrality (30°C) to eliminate basal activity of the muscles or BAT. Consistent with our cold stress experiments, we observed in two different cohorts, a significant lower O_2_ consumption in NCLX KO mice after β-3 stimulation as compared to their WT mice, pointing to a thermogenic dysfunction of BAT in these mice **(Fig. 3d-f and Supplementary Fig. 2c-e)**.

To determine the gene-dose dependent effect we proceeded with evaluation of the heterozygous deletion of NCLX (NCLX +/-). Thermogenic capacity of NCLX +/- mice was also reduced in comparison to their WT littermates, exhibiting significantly lower survival rates and core body temperatures after 6-8 h of cold-stress compared to their WT littermates **(Fig. 3g,h)**.

Furthermore, BAT specific adrenergic activation induced by CL-326,243 injection promoted lower oxygen consumption rates in NCLX +/- mice as compared to WT (3232.5±332 and 2536.7±181 ml/kg/h for WT and NCLX +/- respectively), These results strengthen our hypothesis that the NCLX activity is indispensable for BAT thermogenic function **(Fig. 3i,j)**. Moreover, to determine if the phenotype of NCLX KO mice is gender dependent, we tested our *in vivo* results in female littermates of WT, NCLX +/- and NCLX KO. Both NCLX +/- and NCLX KO female mice are cold-intolerant and have a worse survival rate compared to their WT littermates. **(Supplementary Fig. 3a,b)**.

Overall, these results indicate that mitochondrial Na^+^/Ca^2+^ exchange is essential for adrenergic uncoupled respiration and activation of thermogenesis in BAT.

### Bioenergetics defects in NCLX-null BA are acutely induced following adrenergic stimulation

To investigate the mechanism underlying the suppression of NE-induced uncoupled respiration triggered by NCLX loss and demonstrated *in vitro* and *in vivo*, we first looked for changes in UCP1 expression levels, However we were unable to detect any changes in the protein expression levels in NCLX KO BA as compared to WT BA **(Fig. 4a,b)**. Since a reduction in mitochondrial mass can contribute to the loss of respiratory capacity, we assessed mitochondrial mass by two independent approaches: (1) Western blot analysis of the mitochondrial protein importer TOM20 and (2) Staining with Mitotracker Green, a fluorescent dye that covalently binds to the mitochondria independently of its membrane potential. These experiments showed no indicative changes in mitochondrial mass **(Fig. 4c-e)**.

**Figure 4:**
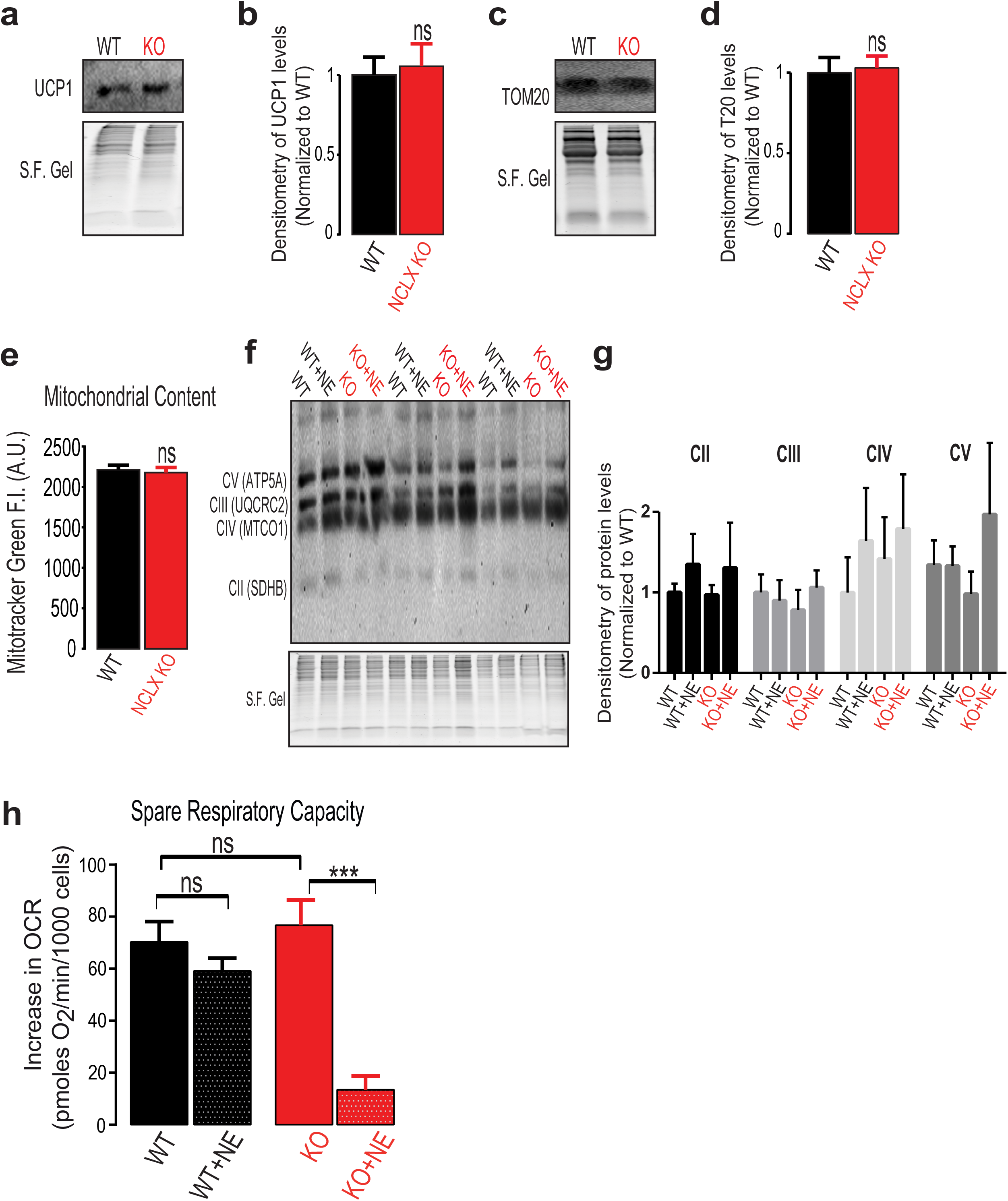
Respiratory dysfunction mediated by NCLX loss in activated BA does not involve changes in UCP1 or mitochondrial mass and is acutely induced following adrenergic stimulation. **(a,b):** Western blot analysis of NCLX-null and WT BA for UCP1 expression levels. **a:** Representative western blots for UCP1 in NCLX-null and WT BA, S.F. Gel was used as a normalization reference for total protein loading. **b:** Quantification of UCP1 levels normalized to S.F. Gel, (n=11 per group **(c,d):** Western blot analysis of NCLX-null and WT BA for TOM20 expression levels. **c:** Representative western blot for TOM20 in NCLX-null and WT BA, S.F. Gel was used for total protein loading. **d:** TOM20 densitometry normalized to S.F. for total protein loading (n=7 per genotype). **e:** Mitotracker Green fluorescence intensity in NCLX-null and WT primary BA, measured by Operetta high-content imaging system. (n=16 per condition from total N=3 imaging experiments). **(f,g):** Western blot analysis of primary NCLX-null and WT BA that were treated with or without NE for 6 h. **f:** Western blot probed with antibodies of OXPHOS complex subunits II–V (CII–CV) for and S.F. Gel was used for total protein loading. **g:** Quantification of each subunit expression of WT and NCLX-null BA with and without NE stimulation normalized to S.F. Gel (n=3-7 per group). **h:** Quantification of maximal spare respiratory capacity in NE stimulated and non-stimulated NCLX-null and WT BA. Maximal respiration was induced by FCCP. Note that spare respiratory capacity is unaltered in non-NE stimulated NCLX KO BA. However, following stimulation with NE, spare respiratory capacity is reduced in the NCLX KO but not in the WT. (N=3-8 independent experiments with n=19-32 per condition). Student’s t-test (**b,d,e**); One-way ANOVA with Tukey’s post-hoc test (**g,h**). Data are expressed as means ± SEM. ns p > 0.05, *** p < 0.0001.

Moreover, Expression levels of OXPHOS complex subunits before and after adrenergic stimulation of BA from NCLX KO and WT were unchanged **(Fig. 4f,g)**.

Interestingly however, spare respiratory capacity before NE stimulation was unaffected in NCLX KO BA compared to their WT. Yet, NE stimulation of NCLX KO led acutely to a collapse of bioenergetics and disability to meet the energy demand **(Fig. 4h)**.

Overall, these results indicate that the mechanism underlying the bioenergetics defect triggered by NCLX loss is functional and acutely induced by NE and independent of UCP1, mitochondrial mass or OXHPOS expression.

### NCLX is essential for maintaining BAT functionality and viability during the activation of thermogenesis

Mitochondrial Ca^2+^ overload can impair respiratory capacity by the induction of permeability transition and mPTP opening. Activation of mPTP involves mitochondrial swelling, release of Cytochrome *c* and can eventually lead to cellular death^8,14,25–27^. We hypothesized that Ca^2+^ overload, induced by NE stimulation in NCLX-deficient BA may induce permeability transition leading to suppression of thermogenesis.

To test this scenario, we employed complementary *in vitro* and *in vivo* models. First, we imaged NCLX KO and WT BA before and after 6 h of NE stimulation. Cells were stained with TOM20 to mark mitochondrial network at the outer membranes and for Cytochrome *c*.

Super resolution confocal microscopy of BA reveals a NE-induced release of Cytochrome *c* from the mitochondria of the NCLX deficient cells but not the WT controls **(Fig. 5a,b)**. In addition, loss of mitochondrial Cytochrome *c* was accompanied by a dramatic mitochondrial swelling and mitochondrial rupture of the outer membrane occurring only under adrenergic stimulation of NCLX-deficient cells **(Fig. 5a,c and Supplementary Fig. 4a,b)**.

**Figure 5:**
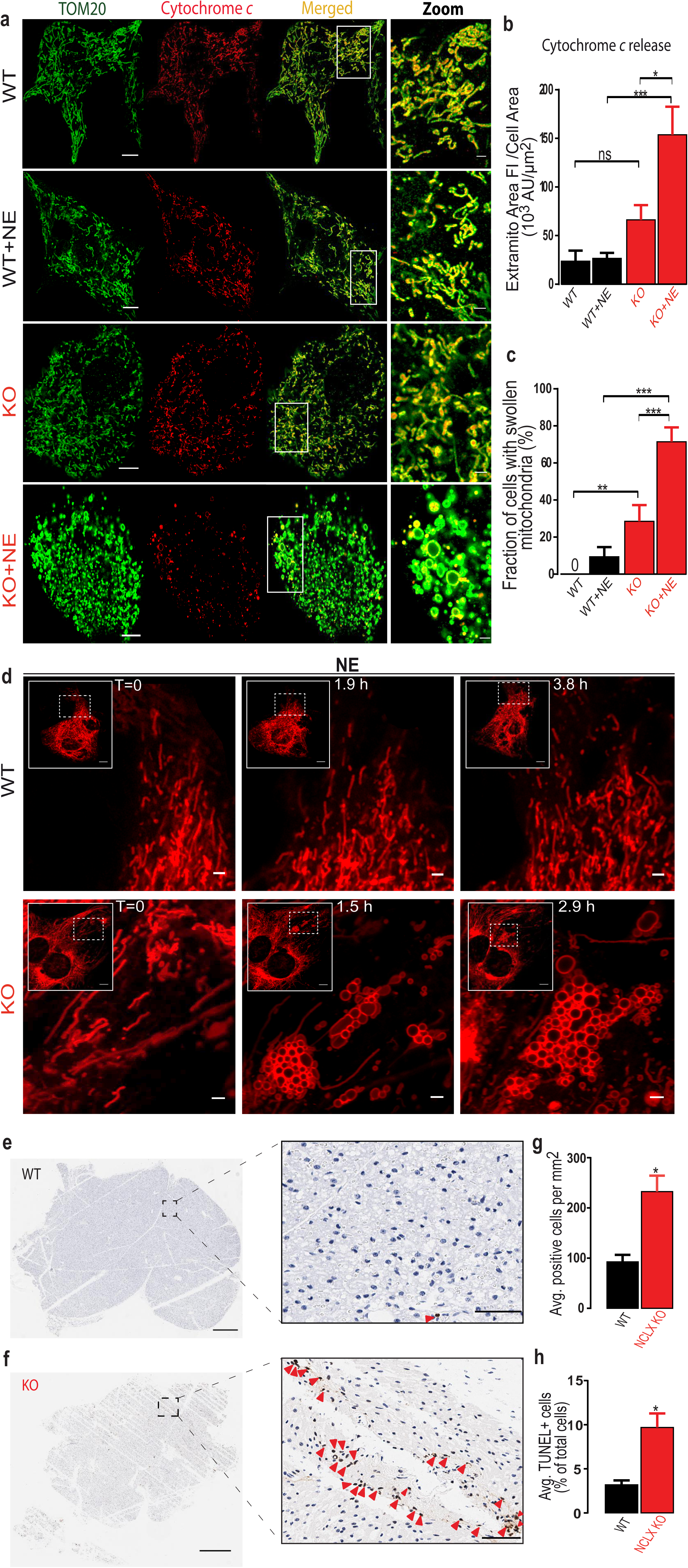
Adrenergic stimulation in NCLX-null BA leads to mitochondrial swelling, cytochrome *c* release, and cell death *in vitro* and *in vivo*. **(a-c):** Super-Resolution confocal images of fixed WT or NCLX-null BA, with or without adrenergic stimulation for 6 h (n=27-46 cells for each of the 4 condition). **a:** NCLX KO and WT BA treated with or without NE and immunostained for Cytochrome *c* (Red). Samples were co-stained for TOM20 to label mitochondrial localization. Note that in NCLX KO BA, NE induces a prominent mitochondrial swelling, accompanied by release of Cytochrome *c* and its dilution in the cytosol. scale bar, 10 and 2 μm for the zoom images. **b:** Quantification of Cytochrome *c* release. Cytochrome *c* in the cytosol was calculated by measuring its total integrated F.I. outside the mitochondria per cell, normalized to total cellular area. **c:** Quantification of mitochondrial swelling. Mitochondrial swelling in each cell was assessed as being present or absent. Zero indicates bars where quantification yielded zero. Note that NE induces a noticeable mitochondrial swelling in NCLX KO BA. **d:** Time-lapse swelling formation of NE stimulated NCLX-null and WT BA. Cells were labelled with mCherry-Fis1 probe marking their outer mitochondrial membrane. Note that swelling spatially starts at one pole of the cell and gradually propagated to the center of the cell. scale bar, 10 and 2 μm for the zoom images. **(e-h):** Cellular death measured by TUNEL staining. Mice were submitted to intermittent cold stress protocol for 5 days. BAT was harvested, fixed and stained with TUNEL. Hematoxylin was used as a counter-staining for all nuclei. (n=3-6 mice per genotype). **e,f:** TUNEL staining of BAT lobes from cold-stressed NCLX KO and WT mice. Note that the number of TUNEL-positive cells (brown; indicated with red arrows) is increased in BAT from cold-stressed NCLX KO as compared to WT mice. scale bar, 500 and 50 μm for the zoom images. **f:** Quantification of the average number of positive cells per cross sectional area in each genotype. **g:** Quantification of the TUNEL-positive cells as percentage of total cells. Student’s t-test (**f,g**); One-way ANOVA with Tukey’s post-hoc test (**b,c**). Data are expressed as means ± SEM. ns p > 0.05, * P<0.05, ** P<0.01, *** p < 0.0001.

To interrogate mitochondrial swelling at high temporal and spatial resolution in adrenergically stimulated NCLX-null BA, mitochondrial outer membranes were labeled with mCherry-Fis1 probe. Time-lapsed imaging showed that mitochondrial swelling of NCLX KO occurs in a spatially organized manner that starts at one pole and gradually propagates to the center of the cell **(Fig. 5d)**.

These results strongly suggest that stimulated BA require NCLX activity to prevent mitochondrial swelling and loss of Cytochrome *c*.

The observation of mitochondrial swelling and Cytochrome *c* release, in NE-stimulated NCLX-null cells in culture suggested that cold exposure may lead to cellular death in BAT. To determine the effect of adrenergic stimulation on cell viability *in vivo* we applied an intermittent cold exposure protocol with the goal of allowing prolonged activation of BAT while preventing hypothermic mortality. Animals were cold-stressed at 4°C for 5 days with recovery periods after a 4-6 h of cold-stress till they gained back their core body temperature. On day 5, BAT was harvested and cellular death was assessed by TUNEL staining of BAT. Evaluation of positive cells per mm^2^ **(Fig. 5e-g)** and the percentage of TUNEL+ cells **(Fig. 5e,f,h)** demonstrate a remarkable cellular death rate of BA from cold-stressed NCLX KO mice as compared to WT **(Fig. 5e,f and Supplementary Fig. 5)**.

Overall, these results indicate that NCLX activity, *in vitro* and *in vivo*, is critical for BA survival during adrenergic stimulation by preventing the ensuing mitochondrial Ca^2+^ overload.

### Blockage of mPTP rescues uncoupled-energy expenditure and thermogenesis in NCLX-null BAT

The findings described in Figure 5 showing mitochondrial swelling and Cytochrome *c* release in activated NCLX-null BA, suggest that in the absence of NCLX, stimulation with NE leads to mitochondrial Ca^2+^ overload and to the induction of permeability transition. A major regulator for the mPTP opening is Cyclophilin D (CypD)^28–30^. Inhibitors of CypD such as Cyclosporin A are potent immune-modulators and have off-targets by interfering with Cyclophilin A and with Calcineurin^31^. However, a novel non-immunosuppressive Cyclosporine derivative, named NIM811, has been shown to effectively block Ca^2+^-induced mPTP opening, without exerting inhibitory side-effects on calcineurin^31,32^.

To test the role of mPTP in mediating the bioenergetics phenotype of NCLX-null BA, we tested the effect of NIM811 on BA response to NE. Primary NCLX KO and WT BA were pretreated either with vehicle (DMSO) or NIM811 and subjected to OCR measurements. We observed that respiratory rates during NE stimulation were not affected by NIM811 in BA of WT mice **(Fig. 6b and Supplementary Fig. 6)**. Strikingly however, NIM811 pretreatment fully restored and rescued NE response in NCLX KO BA with rates similar to the WT BA **(Fig. 6a,b)**. Moreover, this effect was followed by a recovery of maximal respiration **(Fig. 6a)**.

**Figure 6:**
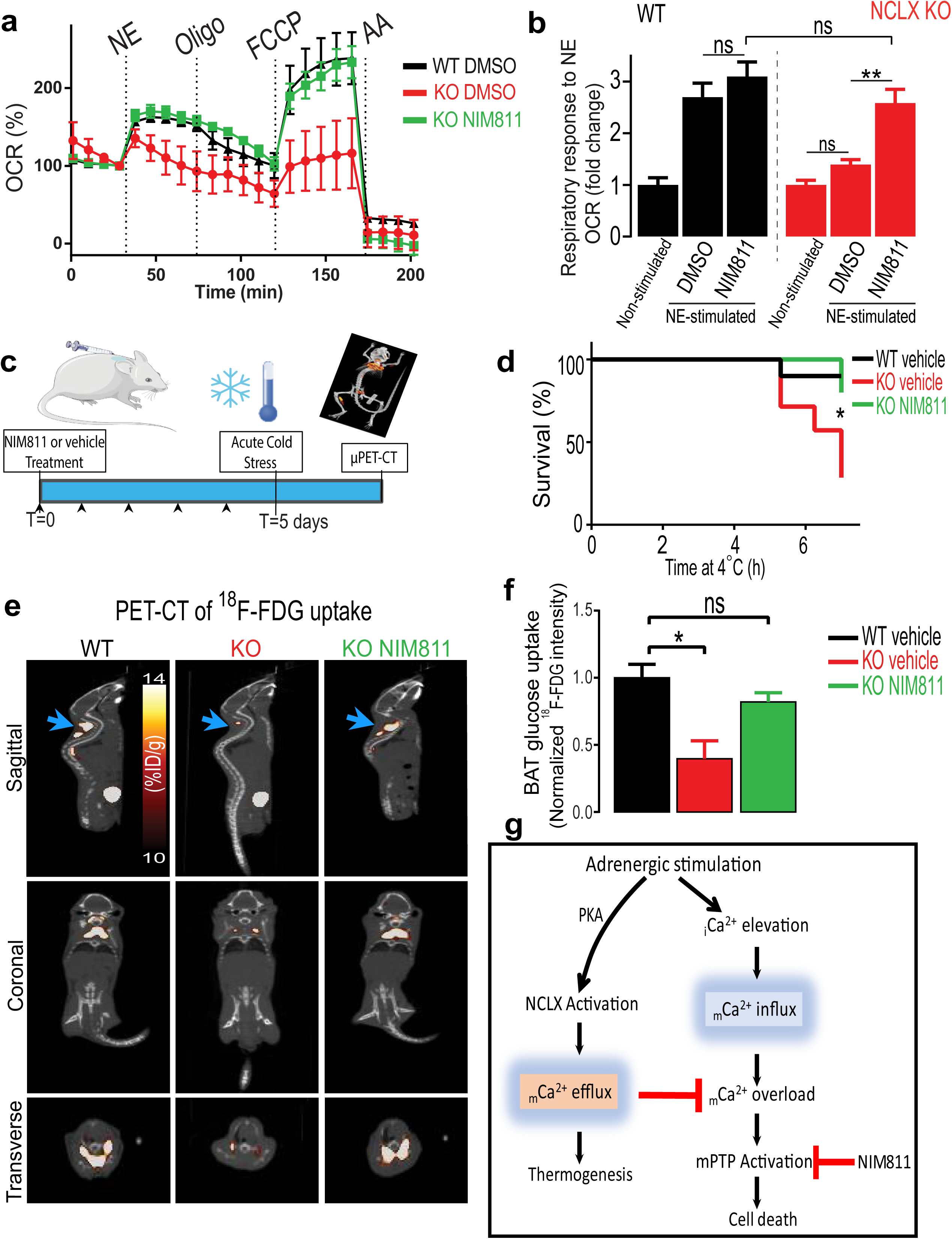
Inhibition of mitochondrial permeability transition in NCLX-null mice restores defective respiratory response to NE *in vitro* and thermogenesis *in vivo*. **(a,b)**: OCR of NCLX-null and WT primary BA that were pretreated either with the mPTP inhibitor, NIM811, or with carrier control, DMSO. Note that NIM811 restored adrenergic stimulated OCR of NCLX-null BA (n=28-36 per condition from N=8-9 independent experiments). **a:** Representative OCR traces in BA from NCLX-KO and WT pretreated either with or without or NIM811 (500nM). **b:** Quantification of NE Response in NCLX KO and WT BA pretreated with or without NIM811. NIM811 rescues respiratory response of NCLX KO BA to NE while the response of WT BA to NE remains unaffected (See Supplementary Fig. 7 for WT treatment traces). **c:** Schematic graph for *in vivo* NIM811experimental procedures. Mice were pretreated for 5 days with a 50 mg/kg dose NIM811 or a vehicle introduced subcutaneously. At the day of experiment mice were acutely cold-stressed (6-8 h) and then immediately PET-imaged with the glucose analog ^18^F-fluorodeoxyglucose (^18^F-FDG) to measure BAT activity under 4°C. **(d,e):** Cold tolerance tests of NCLX KO and WT mice pretreated with or without NIM811 at 4°C with their core body temperature traces and survival curves (n =5-10 mice per group). **d:** Survival curves of 8-10 week old male NCLX KO and WT mice pretreated with or without NIM811 and cold-stressed at 4°C. Mice reaching 28°C or lower were returned to room temperature for recovery. **(e,f):** BAT glucose uptake during cold exposure measured by μPET/CT with ^18^F-FDG (n=3-11 mice per group). **e:** Representative PET-CT images of ^18^F-FDG uptake in cold-stressed mice. Each image is a composition of a CT image superimposed on a PET image. In the CT image, radio-opacity is reflected in grayscale. In the PET image ^18^F-FDG intensity is reflected in hot-metal scale. BAT is marked with white arrows. Note the decreased ^18^F-FDG uptake in NCLX KO BAT, which is completely reversed by NIM811 treatment. ^18^F-FDG uptake is presented as percent injected dose per gram (%ID/g). **f:** Quantification of ^18^F-FDG uptake in BAT of NCLX KO and WT mice pretreated with or without NIM811 as measured by μPET-CT. BAT ^18^F-FDG uptake was normalized to liver uptake as a reference tissue. **g:** Model for the role of NCLX in BAT. NCLX loss of function converts the NE-induced thermogenic pathway to a death pathway due to accumulation of mitochondrial Ca^2+^ and activation of mPTP. NIM811 rescue thermogenesis in NCLX-null BA. intracellular Ca^2+^, _i_Ca^2+^; mitochondrial Ca^2+^,_m_Ca^2+^ One-way ANOVA with Tukey’s post-hoc test (**b,f**); Log-Rank (**d**). Data are expressed as means ± SEM. ns p > 0.05, * P<0.05, ** P<0.01.

We then asked if NIM811 treatment can rescue BAT function of NCLX-null mice *in vivo*. We pretreated NCLX KO mice either with vehicle or NIM811 (50mg*kg^−1^) and compared their thermogenic function to WT mice, this was followed by the measurement of BAT glucose uptake during cold exposure, assessed by μPET/CT imaging with ^18^F-fluorodeoxyglucose (^18^FFDG) analog **(Fig. 6c)**. While cold-stress of 6-8 h exhibited a worse survival rate and impaired cold-tolerance of NCLX KO treated with a vehicle, the NIM811-treated NCLX KO mice displayed an increased overall survival rate, with values remarkably similar to WT animals **(Fig. 6d and Supplementary Fig. 7)**.

We next determined cold-induced BAT activity, assessed by *in vivo* glucose uptake utilizing the ^18^F-FDG analog. While NCLX KO mice showed a reduction of 60.33% ± 18.67% in BAT glucose uptake as compared to WT, the NIM811 pretreated NCLX KO mice almost fully recovered their BAT glucose uptake reaching values similar to those found in WT mice **(Fig. 6e,f and Supplementary Videos 1-3)**.

Overall, this set of data suggests that inhibition of the mPTP by NIM811 can restore NE-induced mitochondrial respiration in NCLX KO BA and rescue thermogenesis in both *in vitro* and *in vivo* systems, thus confirming the role of the mPTP in the pathogenic effect of NCLX deficiency.

## Discussion

Activation of thermogenesis involves cytosolic and mitochondrial calcium elevations, supporting a surge in oxygen consumption and energy expenditure, called upon by adrenergic stimulation of mitochondrial uncoupling and depolarization^5,6,33–36^. Yet, the exact same conditions are shown in other cells to effectively induce permeability transition, mitochondrial swelling and Cytochrome *c* release^8–10^. The mechanism by which brown adipocytes avoid death while responding to a thermogenic signal has not been elucidated. In this study, we have identified a mitochondrial Ca^2+^ extrusion pathway that is activated hormonally by the neurotransmitter NE in BA, and is mediated by the mitochondrial Na^+^/Ca^2+^exchanger, NCLX. Identification of NCLX as the molecular entity regulating mitochondrial Ca^2+^ efflux during thermogenesis allowed us for the first time to look into the mechanism by which BA regulate mitochondrial Ca^2+^ elevation and prevent depolarization-induced permeability transition and death during activation of thermogenesis **(Fig. 6g and Supplementary Fig. 8)**.

*In vitro* and *in vivo* experiments in this study show that stimulation of Ca^2+^extrusion is indispensable for physiological uncoupled energy expenditure in BA. Mechanistically, this is the first study that shows a physiological role of PKA-regulated mitochondrial Ca^2+^ extrusion via NCLX activation. Furthermore, we find that adrenergic activation of NCLX-null BAT impairs mitochondrial bioenergetics through the activation of permeability transition, allowing mitochondrial swelling and Cytochrome *c* release, followed by cell death. Our study and a recent work in cardiac-induced model of NCLX KO^37^ suggest that NCLX acts as an anti-apoptotic mechanism by inhibiting mitochondrial Ca^2+^ overload. Moreover, our study provides a first evidence of a hormonal-stimulus resulting in mitochondrial Ca^2+^ overload and leading to cell death. This supports previous studies demonstrating that, apart from its stimulatory thermogenic role, NE functions as a pro-survival hormone for BAT ^2,38^. In this study we used global NCLX KO, raising the possibility for the involvement of the deletion of NCLX in other tissues. However, there are several reasons supporting that the impaired thermogenesis *in vivo* was due to BAT dysfunction. First, non-shivering assessment by β-3 stimulation and ^18^F-FDG uptake in BAT, are organ-specific measurements in BAT. Second, our *in vitro* system of isolated NCLX KO BA, demonstrating an impaired energy expenditure and remarkable mitochondrial damage, recapitulate the *in vivo* phenotype. These data support the argument that the phenotypes seen in BAT are unlikely to be secondary to other organs.

Remarkably, this study shows that mitochondrial Ca^2+^ overload is not deleterious per se for mitochondrial bioenergetics as long as the mPTP remains closed. Pharmacological inhibition of the mPTP by NIM811 was able to completely rescue the adrenergically-mediated bioenergetics defects and the thermogenic dysfunction of NCLX KO BAT, in *in vitro* as well as *in vivo*. This is in agreement with a recent study showing that NIM811 can prevent cell death induced by mitochondrial Ca^2+^ in liver and mediated by the loss of Micu1, a regulator of MCU ^32^. In summary, the results of this study reveal that NCLX plays an essential role in regulating thermogenic BAT metabolic function and viability. The physiological relevance of PKA-regulation of NCLX and mPTP inhibition shown in the study open new avenues for targeted therapeutic strategies for pathological conditions in which impaired metabolic status is triggered by perturbed mitochondrial Ca^2+^ homeostasis.

## Materials and Methods

### Experimental animals

The mice studies were conducted under an approved Institutional Animal Care and Use Committee (IACUC) protocol at University of California, Los Angeles (UCLA) and Ben-Gurion University. The mice were congenic to the C57BL/6NJ background, fed standard chow diet, and maintained under controlled conditions (housing at 22°C with a 12:12 h light:dark cycle). In all experiments, mice were age and gender matched. For *in vivo* experiments, age-matched male and female mice of 10-12 weeks old were used. For *in vitro* experiments 3-4 weeks old male mice were used.

WT C57BL/6NJ mice and NCLX null (C57BL/6NJ-*Slc8b1^em1(IMPC)J^*/J) were purchased from Jackson laboratories (Jackson lab, Bar Harbor, ME), and bred in our vivarium. Mice genotyping was performed on earpieces or clipped tails obtained during the weaning of pups. Genotyping was performed following the protocol of Jax laboratories by real-time polymerase chain reaction, using a commercial vendor (Transnetyx). The following primers were used to detect NCLX null -/-, Heterozygous +/- or WT +/+ mice:

> Forward Primer-GGCTCCTGTCTTCCTCTGTG and Reverse Primer-GTGTCCATGGGCTTTTGTG.

For *in vivo* experiments and analysis a randomization and a double blinded-manner were performed using ear-tagging and random mice numbering systems which were revealed after the termination of the experiment.

### NIM811 drug preparation and delivery

For mPTP studies, mice were injected subcutaneously above the dorsal brown fat at 50 mg/kg final dose of NIM811 (Novartis). The injection was made once a day for 5 days before cold-stress or μPET/CT experiments. The drug was diluted in a solution containing sterile saline, autoclaved Cremophor EL (15%, v/v) (Kolliphor EL, Sigma, C5135) and sterile Ethanol (5%, v/v) to facilitate administration of NIM811 and its suspension stability.

### Thermogenesis and acute cold exposure

Cold exposure experiments were performed as described previously ^39^. In brief, subcutaneous, biocompatible, and sterile microchip transponders (IPTT-300, Bio Medic Data Systems, Seaford, DE, USA) were implanted in male and female mice in all groups at least 5 days prior to experimentation. On the day of the experiment, mice were housed singly in pre-chilled cages at 4°C with free access to water. Body temperature was assessed every 30-45 min for 6–8 h using a wireless reader system (DAS-8007, Bio Medic Data Systems).

According to our IACUC protocol we defined “survival” when mouse body temperature remained >28°C. At any temperature equal or below 28°C, mice were rescued by removal to room temperature. The time of removal was recorded and considered as drop-out.

For experiments in which cell viability was detected *in vivo*, we applied an intermittent cold exposure which allows for longer term cold exposure while preventing animal death. Animals were exposed to cold at intervals of 4-6 h intermitted by recovery periods till the mice gained back their core body temperature on a course of 5 days.

### *In vivo* measurement of BAT function

In order to evaluate non-shivering thermogenesis and BAT function, we measured whole-body O_2_ consumption at basal and in response to β3-adrenergic agonist, CL316-243 (Sigma, C5976), in anesthetized mice as described previously^40^. Briefly, mice were housed at thermoneutrality (30°C) to eliminate any basal effect of brown fat and muscle shivering activity, mice were anesthetized using pentobarbital (120 mg/kg) at 30°C and placed in metabolic cages with environmental enclosures at 30°C (CLAMS-ENC). After 45-60 min, CL was subcutaneously injected (1 mg/kg) and mice were placed back in the metabolic cages for O_2_ consumption measurements.

To assess BAT function during cold-stress condition, the metabolic cages were pre-chilled at 4°C, mice were again singly housed in the 4°C cages and O_2_ was monitored for 7-8 h.

### PET/CT Imaging

Following 6-8 h of cold-stress, *in vivo* μPET/CT imaging was performed at the Crump Institute Preclinical Imaging Technology Center. Mice were injected with the radioactive glucose analog ^18^F-FDG as reported previously^41^. In brief, mice were injected via lateral tail vein with 70 µCi of ^18^F-FDG followed by 60 min of unconscious uptake under 1.5% isoflurane of anesthesia and cold-conditions. This was followed by a static μPET/CT imaging. a Region-Of-Interest analysis was conducted using AMIDE software^42^ on dorsal BAT. The uptake was normalized to liver as a reference tissue.

### Cell culture

Primary BA were generated by differentiating pre-adipocytes isolated from BAT as described in detail previously^43–45^. BAT was harvested from 3 to 4-weeks-old WT and NCLX KO mice. The tissue was dissected from interscapular, subscapular, and cervical regions, minced, and transferred to a collagenase digestion buffer (2 mg/mL Collagenase Type II in 100 mM HEPES, 120 mM NaCl, 4.8 mM KCl, 1 mM CaCl_2_, 4.5 mM Glucose, 1.5% BSA, pH 7.4) at 37°C under constant agitation for 30 min. Collagenase digestion was performed in 37°C water incubator under constant agitation for 25 min with vortex agitation every 5 min. Digested tissue was homogenized and strained through 100 mm and 40 mm strainers. Cold DMEM was added to tissue digest and centrifuged twice (the last included washing and resuspension in new DMEM at 200 x g speed for 12 min at 4°C). Finally, cell pellets (preadipocytes) were re-suspended 5 mL growth medium (DMEM supplemented with 20% newborn calf serum (NCS), 4 mM Glutamine, 10 mM HEPES, 0.1 mg/mL sodium ascorbate, 50 U/mL penicillin, 50 mg/mL streptomycin) and plated in 6-well plates (Corning). Cells were incubated in 37°C 8% CO_2_ incubator. 24 h after isolation, the cells were washed to remove debris and medium was replaced. 72 h after isolation the cells were lifted using STEMPro Accutase, counted, and re-plated in differentiation media (growth media supplemented with 1 mM rosiglitazone maleate and 4 nM human recombinant insulin). Cells were differentiated for 7 days and medium was changed every other day. For transduction experiments, cells were transduced with virus in differentiation day 0-3 (see below).

For mPTP inhibition experiments, cells at the beginning of differentiation process were co-treated with DMSO, or CyPD inhibitor NIM811 at a concentration of 500nM (Novartis).

### Virus preparation

mCherry-GFP-FIS1(101-152) construct was a generous gift from Ian Ganley^46^, plasmid was packaged into adenoviral particles (Welgen) and was used to stain outer-mitochondrial membrane. For the studies presented here, only the mCherry fluorophore was excited and recorded.

### Measurement of Oxygen Consumption rate

Ten thousand primary brown pre-adipocytes were plated, grown and differentiated for 7 days on a Seahorse 24-well microplate (Agilent, Santa Clara, CA). Oxygen consumption rate of the cells was measured using Seahorse XFe24 Analyzer, as previously described^44,47^. Normalization to cell number was done using the Operetta fluorescence microscope by counting the number of nuclei using Hoechst staining (2 μg/mL, Thermo).

Before the experiment, cells medium was replaced to assay media (DMEM modified medium without sodium bicarbonate (D5030, Sigma) with an addition of 3 mM glucose and 2 mM Glutamine, followed by incubation for 60 min at 37°C (in a non-CO_2_ incubator) before loading into the XFe24 extracellular analyzer (Agilent). During these 60 min the ports of the cartridge containing the oxygen probes were loaded with the compounds to be injected during the assay (50 □L/port) and the cartridge was calibrated.

After steady basal consumption rates were obtained, NE was injected first to a final concentration of 1.5 μM. This was followed by injection of Oligomycin (Calbiochem, San Diego, CA, US) at a final concentration of 2 μM, followed by injection of (carbonyl cyanide 4-(trifluoromethoxy) phenylhydrazone) FCCP (Sigma, C2920) at a final concentration of 2 μM, Finally, 3 μM Antimycin A (sigma, A8674) was injected to get non-mitochondrial respiration fraction.

### siRNA preparation and transfection

Double-stranded siRNAs used to silence NCLX expression was obtained from Ambion (Applied Biosystems) as reported before ^17^. The sequence of 21 nucleotides corresponding to the sense strands used for the NCLX siRNA was AACGGCCACUCAACUGUCUtt and that for the control siRNA was AACGCGCAUCCAACUGUCUtt. siNCLX or siControl were diluted in Lipofectamine 3000 transfection reagent according to the manufacturer’s instructions (Thermo). The efficiency of transfection was assessed by visualizing co-transfecting Dharmacon siGLO Green transfection particles according to the protocol provided by manufacturer (Dharmacon, D-001630-01-05). The transfection efficiency, for siNCLX delivery as determined by siGlo fluorescent marker was ~70-90%.

### High-throughput Imaging

Cells stained with Mitotracker Green (MTG) were imaged on PerkinElmer Operetta high-content wide-field fluorescence imaging system, coupled to Columbus analysis software. 40× NA objective lens was used, in a single focal plane across each plate. For long-term imaging experiments, the cells kept inside an environmental chamber at 37°C and 5% CO_2_ levels. The bottom of each well was detected automatically by the Operetta focusing laser, and the focal plane calculated relative to this value. MTG excitation (460-490 nm) emission (520-550 nm) was imaged for ~100 ms with a total of 15 fields of view taken per well, with an identical pattern of fields used for every well.

Modified Columbus (PerkinElmer) image analysis software was used to calculate intensity of the fluorescent probe.

### Fluorescent Ca^2+^ imaging

Kinetic live-cell fluorescent imaging was performed to monitor Ca^2+^ transients using an imaging apparatus that consisted of an Axiovert 100 inverted microscope (Zeiss, Oberaue, Germany), Polychrome V monochromator (Till Photonics, Planegg, Germany) and a Sensi-Cam cooled charge-coupled device (PCO, Kelheim, Germany). Fluorescence images were acquired with Imaging WorkBench 6.0 software (Axon Instruments, Foster City, CA, USA). Ca^2+^ imaging was performed in BA that were grown and attached onto coverslips and superfused with a Krebs-Ringer’s solution containing (in mM): 123 NaCl, 5.4 KCl, 0.8 MgCl_2_, 20 HEPES, 1.8 CaCl_2_, 15 D-Glucose, 2 Glutamine and 1% free-fatty acid Bovine Serum Albumin (BSA).; pH was adjusted to 7.4 with NaOH. In some experiments, NMDG^+^ was used to replace NaCl in Na^+^-free Krebs-Ringer’s solutions and pH was adjusted to 7.4 with KCl^17^. For mitochondrial Ca^2+^ measurements, cells were washed and then loaded with Rhod-2 AM (1 μM) for 30 min at 37°C. After loading cells were washed again three times followed by additional incubation of 30 min to allow for the de-esterification of the dye. Rhod-2AM was excited at 552 nm wavelength light and imaged with a 570 nm long-pass filter.

Mitochondrial Ca^2+^ response was triggered by switching the perfusion solution to Ca^2+^ – free Ringer’s solution supplemented by NE (1.5 μM) or ATP (100 μM). In some experiments, cells were pretreated with Forskolin (Sigma-Aldrich) or H-89 (Santa Cruz Biotechnology), at 50μM and 5μM concentrations, respectively.

Traces of Ca^2+^ responses were analyzed and plotted using KaleidaGraph. The rate of ion transport was calculated from each graph (summarizing an individual experiment) by a linear fit of the change in the fluorescence (ΔF) for Ca^2+^ influx and efflux over time (ΔF/dt). Rates from n experiments (as mentioned in legends to the figures) were averaged and displayed in bar graph.

### Super Resolution Microscopy

A Zeiss LSM 880 confocal microscope with Airyscan mode was used for super-resolution imaging, with 488 nm and 561nm lasers and 63×objective. 488 nm laser was used to excite Alexa Fluor 488 while 561 nm laser was used for excitation of Alexa Fluor 568 and mCherry fluorophore (in experiments of BA infected with mCherry-GFP-Fis-1 AAV). At least 25 cells per condition were collected, the cells were individualized as areas of interest using FIJI ImageJ software (https://fiji.sc/), and Further image analysis was applied on them as described below.

### Fluorescent dyes

Mitotracker Green (MTG), (Thermo) was used at 200 nM. Staining proceeded for 45 min followed by 3 washes with PBS at 37°C. before imaging. Rhod-2 AM (Thermo) was used at 1 μM, cells were stained for 30 min at 37°C. Dye was washed three times with Ringer’s buffer and re-incubated for an extra 30 min to allow de-esterification of the intracellular AM esters. Hoechst was used at 2 μg/mL concentration (Sigma, 33258) to stain the nuclei for 30 min prior to Operetta experiments for counting nuclei of Seahorse assays.

### Western Blotting and antibodies

After treatment, cells were rinsed three times with ice-cold PBS and scraped in an ice-cold RIPA lysis buffer (containing 50 mmol/liter Tris-HCl, pH 7.5, 0.1% (w/v) Triton X-100, 1 mmol/liter EDTA, 1 mmol/liter EGTA, 50 mmol/liter NaF, 10 mmol/liter sodium β-glycerophosphate, 5 mmol/liter sodium pyrophosphate, 1 mM sodium vanadate, and 0.1% (v/v) 2-mercaptoethanol) and protease inhibitors (a 1:1,000 dilution of protease inhibitor mixture; Sigma P8340). The lysates were shaken for 20 min at 4°C, centrifuged (13,500 x g, 15 min at 4°C), and the supernatant was collected. Protein concentration was determined using the BCA protein assay (Pierce). Equal amounts of protein (12 μg) were mixed with 4X LDS sample buffer before running on 15% SDS-polyacrylamide gel electrophoresis and transferred onto a polyvinylidene difluoride membrane (GVS Life Sciences, #1214429) using a wet transfer system (BioRad Hercules, California, US). The membranes were blocked with 5% nonfat dry milk for 1 h and then incubated with The following antibodies: Total OXPHOS Rodent WB Antibody Cocktail (1:1000, Abcam, ab110413), TOMM20 (1:2000, Santa Cruz, Sc-11415), UCP1 (1:1000, Abcam, ab10983), NCLX (1:1000, Santa Cruz SC-161921 and SC-161922), MCU (1:1000, Santa Cruz, SC-515930).

Antibodies were used according to the manufacturer’s instructions. After overnight incubation, membranes were washed with phosphate-buffered saline containing 0.1% Tween-20 and then incubated with anti–rabbit IgG secondary antibody (Cell Signaling Technology) solution (1:5,000) for 1 h or anti-mouse (1:2,000) for 1 h. The membrane was again washed as above and then exposed to a chemiluminescent protein detection system (ChemiDoc, MP Imaging system, Bio-Rad).

As an alternative for beta actin, stain-free blotting was used as a normalization method according to manufacturer’s instructions (Bio-Rad). Briefly, 0.6% Trichloroethanol (Sigma, T54801) was added to the separating gel. At the end of the electrophoresis of the gel, the gel was exposed to UV for 1 min in order to conjugate tryptophans of the samples with the TCE. After the transfer, the membrane was taken to the ChemiDoc MP system, and an image of the marked proteins was taken. The normalization factor was calculated using the Image lab software, version 5.2.1 (Bio-Rad, Hercules, California, US).

### Immunostaining and Immunocytofluorescence

Cells were cultured and differentiated on quadrant dishes and fixed at 4% vol/vol Paraformaldehyde (PFA) for 15 min at room temperature. After washing three times in PBS, cells were incubated in permeabilization buffer (2 L/mL Triton X-100 and 0.5mg/mL Sodium Deoxycholate in PBS, pH 7.4) for 15 min at room temperature. Subsequently, cells were blocked with 3% BSA for 1 h at room temperature. Next, cells were incubated with 1:200 primary antibody of Cytochrome *c* (Abcam, ab110325) at 4°C overnight. The next day, cells were washed in PBS and incubated with 1:200 primary antibody of TOMM20 (Tom20 FL-145, Santa Cruz) at 4°C for overnight. On the last day, 1:500 Anti-Rabbit Alexa Fluor or 488 (A11008, Thermo) or Anti-Mouse Alexa Fluor 568 antibodies (Thermo, A11004) for 1 h at room temperature. samples were kept in PBS.

### Image Analysis

Cytochrome *c* release was determined by quantifying integrated extra-mitochondrial FI (AU) using FIJI software and normalized per total cell area (μm^2^). Swelling was assessed by the following criteria: visual inspection for fragmented and spherical mitochondria that have a diameter > 4 μm. Cells with more than 5 mitochondria that were positive for these criteria were scored positive.

### Preparation of images for print

Representative microscopy images were transformed from the microscopy format generated by Zen Software (.CZI) to .TIF images using FIJI software. Images were cropped for detail, separated into respective channels, and the *window* and *level* parameters were adjusted identically per channel in all images to emphasize the fluorescent structures in the images without manipulation of raw pixel values. Images were then inserted in PowerPoint software and photo-correction was used to enhance the brightness of all images by increasing it by 40% and lowering the contrast by 40% for illustrative purposes.

Western blot images were also transferred to PowerPoint. When Brightness/Contrast were applied, it was done equally to control and experimental groups including all blot area. For the mCherry-Fis1 experiments, the aim is to show morphological changes. Therefore, Brightness and contrast for each condition were optimized using FIJI taking into account the image’s histogram (*window* and *level* parameters were not identical).

### Histology and Immunohistochemistry Analysis

For *in vivo* cell death assessment in BAT, mice were continuously cold-stressed for 4 days, 6-8 h each day. On the last day, mice were sacrificed and BAT was excised and fixed overnight for 16 h in 10% buffered formalin and then placed in 70% ethanol. Tissues were then processed and embedded by Translational Pathology Core Laboratory (TPCL) at UCLA and TUNEL ApopTag staining was performed on them according to the following the kit instructions (Millipore, S7101), hematoxylin was used for counterstaining. Slides were then scanned onto an Aperio ScanScope AT at 20X magnification (Aperio Technologies, Inc., Vista, CA). Digital slides were blindly analyzed with QuPath software in order to determine percent and area of TUNEL positive cells^48^.

### Statistics

All data are expressed as mean ± SEM. Statistical analysis was performed using Prism 6.0 (GraphPad Software) and two-tailed Student’s *t*-test was used for two groups, Two-way ANOVA followed by Sidak’s test was used for multiple comparisons involving two groups and one-way ANOVA with post-hoc Tukey’s test was used for multiple comparisons.

For *in vivo* studies including μPET/CT imaging, histological sectioning, staining and analysis, investigators were blinded from mouse genotype and an ear tagging system enabled unbiased data collection. *In vivo* experiments involved core facilities that were not familiar with the study, limiting biased results.

## Additional Information

### Data Availability

Source data including imaging, blots data are available on reasonable request.

### Code Availability

Macros built for quantification of Cytochrome *c* release is available on reasonable request.

## Acknowledgemnts

The authors would like to thank those who contributed helpful discussions, insight and support of the research, including Drs. Kiana Mahdaviani, Ilan Benador, Eleni Ritou, Evan Taddeo, Rebeca Acin-Perez, Linsey Stiles, Nour Alsabeeh, Anton Petcherski, Fernanda Cerqueirra, Kyle Trudeau from the Shirihai lab and Drs. Marko Kostic and Soumtira Roy from Sekler lab. The authors would like to thank Drs. Assaf Rudich, Alicia Kowaltowski, Dani Dagan, Barbara Corkey, Marc Prentki, Michal Hershfinkel, Ajit Divakaruni, Ehud Ohana and Daniel Khananshvili for their helpful advices and discussions.

We thank Dr. Jason T. Lee and Charles Zamilpa at UCLA’s Crump Imaging Technology Center for assistance with PET/CT imaging of the mice. We thank Translational Pathology Core Laboratory at UCLA’s DGSOM for assistance with histology samples preparation and processing.

E.A. A. was supported by the Kreitman predoctoral scholarship from Ben-Gurion University and the Israeli Council for higher Education fellowship. O.S. S. is funded by National Institutes of Health grants RO1 DK35914, R01 DK56690, and R01 DK074778. I.S. is funded by Israel Science Foundation (ISF, 1424/17), ISF-China (1210/14) and German-Israeli Project Cooperation (DIP, SE2372/1-1). M.L. is funded by DRC UCLA/UCSD Pilot grant (NIH P30 DK063491) and the Department of Medicine Chair at UCLA.

## Author Contributions

E.A.A. designed and performed experiments, analyzed the data and wrote the manuscript. O.S.S. and I.S. helped design the study and supervised manuscript writing. A.E.J. designed and performed mouse experiments and analyzed PET/CT data. M.V. helped managing the mice colony. M.T. performed BAT isolation and culture. N.A.M. built image analysis macros and helped with microscopy. M.S. helped operating the metabolic cages. G.L., M.F.O. helped with data interpretation and manuscript editing. M.L. helped with experiments design, data interpretation and manuscript editing. All authors read and approved the final version of the manuscript.

## SUPPLEMENTARY FIGURES

**Supplementary Figure 1, related to Figure 2:** Lack of Na^+^ impairs NE-stimulated respiration in BA.

**a:** Representative OCR of BA incubated either with or without Na^+^ (NMDG^+^ is used as a cationic replacement) normalized to cell number. Note that in the addition to the effect seen in NCLX KO BA (Decreased NE and FCCP responses), basal respiration was lower, may be due to off targets of Na^+^ loss.

**b:** Representative OCR traces normalized to basal respiration before NE addition.

**c,d:** Quantification of NE response and spare respiratory capacity after NE stimulation of BA incubated with or without Na^+^. Note that NE-induced uncoupled respiration was suppressed in absence of Na^+^ as well and accompanied by loss of spare respiratory capacity demonstrated by the inability of FCCP to further uncouple the mitochondria, supporting that Na^+^-mediated-Ca^2+^ efflux by mitochondrial Na^+^/Ca^2+^ exchange is essential for uncoupled respiration. N=3 independent experiments with n=18-19 total wells per condition.

Student’s t-test (**c,d**). Data are expressed as means ± SEM. * P<0.05, ** P<0.01, *** p < 0.0001.

**Supplementary Figure 2, Related to Figure 3**

**(a,b):** Cold tolerance tests at 4°C of NCLX KO and WT male littermates with their raw core body temperature traces and survival curves (n =4-6 mice per group).

**a:** Survival curves of 8-10 week old male NCLX KO and WT mice cold-stressed at 4°C. Mice reaching 28°C or lower were returned to room temperature for recovery.

**b:** Core body temperature traces of the animals from the experiment shown in (a) during cold exposure (4°C). Note that the NCLX KO animals display impaired thermogenic capacity as compared to WT. dashed red line indicates animal removal.

**(c-e):** VO2 of male littermates of NCLX KO and WT mice at basal and after CL316, 243-injection (1 mg/kg), under anesthesia at 30°C. (n = 4-6 mice per group).

**c:** Traces of VO2 of NCLX KO and WT mice.

**d:** VO2 at baseline and under CL-stimulation.

**e:** Fold increase of *O_2_* consumption after the CL-stimulation. Note that NCLX KO mice have reduced ability to increase their BAT stimulated *O_2_* consumption compared to WT.

Student’s t-test (**d,e**); Two-way ANOVA (**c**); Log-Rank statistics (**a**). Data are expressed as means ± SEM. *** p < 0.0001.

**Supplementary Figure 3, Related to Figure 3**

**(a,b):** Cold tolerance tests (4°C) of WT, NCLX (+/-) and NCLX KO female littermates with their raw core body temperature traces and survival curves (n =4-5 mice per group).

**a:** Survival curves of 8-10 week old females of WT, NCLX (+/-) and NCLX KO mice cold-stressed at 4°C. Mice reaching 28°C or lower were returned to room temperature for recovery.

**b:** Core body temperature traces of the animals from the same experiment in (A) during cold exposure (4°C).

**Supplementary Figure 4, Related to Figure 5.**

**(a,b):** Super-resolution confocal images for NE-stimulated NCLX KO BA, co-immunostained for TOM20 for marking mitochondrial network (Green) and for Cytochrome *c* (Red). Note that in NCLX KO BA, in addition to mitochondrial swelling and cytochrome *c* loss, NE induces morphological changes to mitochondria including the rupture of the mitochondrial membranes as indicated by the white arrows.

**Supplementary Figure 5, Related to Figure 5.**

Representative of the quantification method of TUNEL positive cells in histology. Red circles indicate positive cells and Blue circles indicate negative cells.

**Supplementary Figure 6, Related to Figure 6.**

Representative OCR in response to NE in BA from WT. Cells were pretreated either with DMSO (control) or NIM811 (500nM). Note that NIM811 does not alter respiration in WT BA.

**Supplementary Figure 7, Related to Figure 6.**

**a:** Core body temperature traces during cold exposure (4°C) of the animals from the experiment shown in Figure 6a. Note that the NCLX KO animals display impaired thermogenic capacity as compared to WT, however NCLX KO mice pretreated with NIM811 were protected. Dashed red line indicates animal removal from 4°C.

**Supplementary Figure** 8

A graphical abstract for the role of NCLX under adrenergic activation of BAT. Full description in Figure 6g and the Discussion section.

## SUPPLEMENTARY VIDEOS

**Supplementary Video 1, Related to Figure 6.**

Video of ^18^F-FDG-μPET/CT Imaging of cold-induced glucose uptake in a representative WT mouse.

**Supplementary Video 2, Related to Figure 6.**

Video of ^18^F-FDG-μPET/CT Imaging of cold-induced glucose uptake in a representative NCLX KO mouse.

**Supplementary Video 3, Related to Figure 6.**

Video of ^18^F-FDG-μPET/CT Imaging of cold-induced glucose uptake in a representative NCLX KO mouse pretreated with NIM811.

## References

1. Nedergaard, J. & Cannon, B. The changed metabolic world with human brown adipose tissue: therapeutic visions. Cell Metab. 11, 268–272 (2010).

2. Cannon, B. & Nedergaard, J. Brown adipose tissue: function and physiological significance. Physiol. Rev. 84, 277–359 (2004).

3. Divakaruni, A. S. & Brand, M. D. The regulation and physiology of mitochondrial proton leak. Physiology (Bethesda). 26, 192–205 (2011).

4. Nicholls, D. G. The physiological regulation of uncoupling proteins. Biochim. Biophys. Acta 1757, 459–466 (2006).

5. Territo, P. R., French, S. A., Dunleavy, M. C., Evans, F. J. & Balaban, R. S. Calcium activation of heart mitochondrial oxidative phosphorylation: rapid kinetics of mVO2, NADH, AND light scattering. J. Biol. Chem. 276, 2586–2599 (2001).

6. Denton, R. M. Regulation of mitochondrial dehydrogenases by calcium ions. Biochim. Biophys. Acta 1787, 1309–1316 (2009).

7. Drago, I., Pizzo, P. & Pozzan, T. After half a century mitochondrial calcium in- and efflux machineries reveal themselves. EMBO J. 30, 4119–4125 (2011).

8. Kwong, J. Q. & Molkentin, J. D. Physiological and pathological roles of the mitochondrial permeability transition pore in the heart. Cell Metab. 21, 206–214 (2015).

9. Nicholls, D. G. Mitochondrial calcium function and dysfunction in the central nervous system. Biochim. Biophys. Acta 1787, 1416–1424 (2009).

10. Szabó, I. & Zoratti, M. The mitochondrial megachannel is the permeability transition pore. J. Bioenerg. Biomembr. 24, 111–117 (1992).

11. Morciano, G. et al. Molecular identity of the mitochondrial permeability transition pore and its role in ischemia-reperfusion injury. J. Mol. Cell. Cardiol. 78, 142–153 (2015).

12. Di Lisa, F., Menabo, R., Canton, M., Barile, M. & Bernardi, P. Opening of the mitochondrial permeability transition pore causes depletion of mitochondrial and cytosolic NAD+ and is a causative event in the death of myocytes in postischemic reperfusion of the heart. J. Biol. Chem. 276, 2571–2575 (2001).

13. Bernardi, P. Modulation of the mitochondrial cyclosporin A-sensitive permeability transition pore by the proton electrochemical gradient. Evidence that the pore can be opened by membrane depolarization. J. Biol. Chem. 267, 8834–8839 (1992).

14. Giorgio, V., Guo, L., Bassot, C., Petronilli, V. & Bernardi, P. Calcium and regulation of the mitochondrial permeability transition. Cell Calcium 70, 56–63 (2018).

15. Baughman, J. M. et al. Integrative genomics identifies MCU as an essential component of the mitochondrial calcium uniporter. Nature 476, 341–345 (2011).

16. De Stefani, D., Raffaello, A., Teardo, E., Szabo, I. & Rizzuto, R. A forty-kilodalton protein of the inner membrane is the mitochondrial calcium uniporter. Nature 476, 336–340 (2011).

17. Palty, R. et al. NCLX is an essential component of mitochondrial Na+/Ca2+ exchange. Proc. Natl. Acad. Sci. U. S. A. 107, 436–441 (2010).

18. Gunter, T. E. & Pfeiffer, D. R. Mechanisms by which mitochondria transport calcium. Am. J. Physiol. 258, C755-86 (1990).

19. Al-Shaikhaly, M. H., Nedergaard, J. & Cannon, B. Sodium-induced calcium release from mitochondria in brown adipose tissue. Proc. Natl. Acad. Sci. U. S. A. 76, 2350–2353 (1979).

20. Kostic, M. et al. PKA Phosphorylation of NCLX Reverses Mitochondrial Calcium Overload and Depolarization, Promoting Survival of PINK1-Deficient Dopaminergic Neurons. Cell Rep. 13, 376–386 (2015).

21. Bagur, R. & Hajnoczky, G. Intracellular Ca(2+) Sensing: Its Role in Calcium Homeostasis and Signaling. Mol. Cell 66, 780–788 (2017).

22. White, N. & Burnstock, G. P2 receptors and cancer. Trends Pharmacol. Sci. 27, 211–217 (2006).

23. Koshimizu, T. A. et al. Characterization of calcium signaling by purinergic receptor-channels expressed in excitable cells. Mol. Pharmacol. 58, 936–945 (2000).

24. Berridge, M. J. Inositol trisphosphate and calcium signalling. Nature 361, 315–325 (1993).

25. Giorgio, V. et al. Ca(2+) binding to F-ATP synthase beta subunit triggers the mitochondrial permeability transition. EMBO Rep. 18, 1065–1076 (2017).

26. Giorgio, V. et al. Dimers of mitochondrial ATP synthase form the permeability transition pore. Proc. Natl. Acad. Sci. U. S. A. 110, 5887–5892 (2013).

27. Alavian, K. N. et al. An uncoupling channel within the c-subunit ring of the F1FO ATP synthase is the mitochondrial permeability transition pore. Proc. Natl. Acad. Sci. U. S. A. 111, 10580–10585 (2014).

28. Basso, E. et al. Properties of the permeability transition pore in mitochondria devoid of Cyclophilin D. J. Biol. Chem. 280, 18558–18561 (2005).

29. Baines, C. P. et al. Loss of cyclophilin D reveals a critical role for mitochondrial permeability transition in cell death. Nature 434, 658–662 (2005).

30. Elrod, J. W. et al. Cyclophilin D controls mitochondrial pore-dependent Ca(2+) exchange, metabolic flexibility, and propensity for heart failure in mice. J. Clin. Invest. 120, 3680–3687 (2010).

31. Waldmeier, P. C., Feldtrauer, J.-J., Qian, T. & Lemasters, J. J. Inhibition of the mitochondrial permeability transition by the nonimmunosuppressive cyclosporin derivative NIM811. Mol. Pharmacol. 62, 22–29 (2002).

32. Antony, A. N. et al. MICU1 regulation of mitochondrial Ca(2+) uptake dictates survival and tissue regeneration. Nat. Commun. 7, 10955 (2016).

33. Ye, L. et al. TRPV4 is a regulator of adipose oxidative metabolism, inflammation, and energy homeostasis. Cell 151, 96–110 (2012).

34. Chen, Y. et al. Crosstalk between KCNK3-Mediated Ion Current and Adrenergic Signaling Regulates Adipose Thermogenesis and Obesity. Cell 171, 836–848.e13 (2017).

35. Sun, W. et al. Lack of TRPV2 impairs thermogenesis in mouse brown adipose tissue. EMBO Rep. 17, 383–399 (2016).

36. Ikeda, K. et al. UCP1-independent signaling involving SERCA2b-mediated calcium cycling regulates beige fat thermogenesis and systemic glucose homeostasis. Nat. Med. 23, 1454–1465 (2017).

37. Luongo, T. S. et al. The mitochondrial Na(+)/Ca(2+) exchanger is essential for Ca(2+) homeostasis and viability. Nature 545, 93–97 (2017).

38. Nedergaard, J., Wang, Y. & Cannon, B. Cell proliferation and anti-apoptosis: Essential processes for recruitment of the full thermogenic capacity of brown adipose tissue. Biochim. Biophys. Acta (2018). doi:10.1016/j.bbalip.2018.06.013

39. Mahdaviani, K. et al. Mfn2 deletion in brown adipose tissue protects from insulin resistance and impairs thermogenesis. EMBO Rep. 18, 1123–1138 (2017).

40. Cannon, B. & Nedergaard, J. Nonshivering thermogenesis and its adequate measurement in metabolic studies. J. Exp. Biol. 214, 242–253 (2011).

41. Momcilovic, M. et al. Targeted Inhibition of EGFR and Glutaminase Induces Metabolic Crisis in EGFR Mutant Lung Cancer. Cell Rep. 18, 601–610 (2017).

42. Loening, A. M. & Gambhir, S. S. AMIDE: a free software tool for multimodality medical image analysis. Mol. Imaging 2, 131–137 (2003).

43. Cannon, B. & Nedergaard, J. Cultures of adipose precursor cells from brown adipose tissue and of clonal brown-adipocyte-like cell lines. Methods Mol. Biol. 155, 213–224 (2001).

44. Wikstrom, J. D. et al. Hormone-induced mitochondrial fission is utilized by brown adipocytes as an amplification pathway for energy expenditure. EMBO J. 33, 418–436 (2014).

45. Benador, I. Y. et al. Mitochondria Bound to Lipid Droplets Have Unique Bioenergetics, Composition, and Dynamics that Support Lipid Droplet Expansion. Cell Metab. 27, 869–885.e6 (2018).

46. Allen, G. F. G., Toth, R., James, J. & Ganley, I. G. Loss of iron triggers PINK1/Parkin-independent mitophagy. EMBO Rep. 14, 1127–1135 (2013).

47. Wu, M. et al. Multiparameter metabolic analysis reveals a close link between attenuated mitochondrial bioenergetic function and enhanced glycolysis dependency in human tumor cells. Am. J. Physiol. Cell Physiol. 292, C125-36 (2007).

48. Bankhead, P. et al. QuPath: Open source software for digital pathology image analysis. Sci. Rep. 7, 16878 (2017).

